# Whole genome sequencing and assembly of a *Caenorhabditis elegans* genome with complex genomic rearrangements using the MinION sequencing device

**DOI:** 10.1101/099143

**Authors:** JR Tyson, NJ O’Neil, M Jain, HE Olsen, P Hieter, TP Snutch

## Abstract

Advances in 3^rd^ generation sequencing have opened new possibilities for ‘benchtop’ whole genome sequencing. The MinION is a portable device that uses nanopore technology and can sequence long DNA molecules. MinION long reads are well suited for sequencing and *de novo* assembly of complex genomes with large repetitive elements. Long reads also facilitate the identification of complex genomic rearrangements such as those observed in tumor genomes. To assess the feasibility of the *de novo* assembly of large complex genomes using both MinION and Illumina platforms, we sequenced the genome of a *Caenorhabditis elegans* strain that contains a complex acetaldehyde-induced rearrangement and a biolistic bombardment-mediated insertion of a GFP containing plasmid. Using ∼5.8 gigabases of MinION sequence data, we were able to assemble a *C. elegans* genome containing 145 contigs (N50 contig length = 1.22 Mb) that covered >99% of the 100,286,401 bp reference genome. In contrast, using ∼8.04 gigabases of Illumina sequence data, we were able to assemble a *C. elegans* genome in 38,645 contigs (N50 contig length = ∼26 kb) containing 117 Mb. From the MinION genome assembly we identified the complex structures of both the acetaldehyde-induced mutation and the biolistic-mediated insertion. To date, this is the largest genome to be assembled exclusively from MinION data and is the first demonstration that the long reads of MinION sequencing can be used for whole genome assembly of large (100 Mb) genomes and the elucidation of complex genomic rearrangements.

## Introduction

Advances in Next Generation Sequencing (NGS) have ushered in a new era of whole genome analysis. The short sequencing reads generated by sequencing-by-synthesis NGS are well suited for resequencing complex genomes for which a reference sequence has been established. However, short reads are poorly suited for *de novo* assembly of complex genomes in part due to repeat regions that generate highly discontinuous assemblies and require unique flanking sequences to position repeat elements. In many instances if a sequencing read does not completely span a repeat, either due to repeat size or physical properties of the repetitive DNA, it can not be unambiguously assembled into a contig. This can pose a problem for the *de novo* assembly of metazoan genomes where repetitive elements are common. For example, approximately 12% of the 100 Mb *C. elegans* genome is derived from transposable elements (Bessereau JL. 2006). As such, NGS is not optimal for sequencing and assembly of large novel genomes or genomes with numerous complex rearrangements such as those observed in tumour cells.

Third Generation Sequencing (TGS) technologies have increased read lengths 100 to 1000-fold compared to NGS platforms and therefore can span much larger repeat regions than NGS. The first major TGS platform was the Pacific Biosciences Single Molecule Real Time (SMRT) PacBio sequencing system. PacBio sequencing averages 10 kb read lengths and with consensus sequencing the error rate approaches that of NGS. In 2014, Oxford Nanopore Technologies (ONT) introduced the MinION nanopore sequencer. The MinION directly connects to a laptop or desktop PC via a USB3 port, is ∼90 grams, and costs are for reagents only. The MinION works on the principle of nanopore strand sequencing (Deamer et al. 2016). Briefly, a DNA strand is enzymatically unzipped and ratcheted through a membrane-inserted protein nanopore as a voltage potential is applied. As the DNA single strand translocates, the resulting changes in ionic current are measured and sequence calls produced corresponding to the identity of the nucleotides via computationally inferred model fitting. When only one strand (template) from a duplex DNA molecule is read by the nanopore, the resulting sequence call is termed a 1D read (Jain et al 2016). With certain library preparations an optional hairpin strand bridge provides the opportunity to sequence both strands of a dsDNA (template and complement) generating a 2D read, which results in a higher accuracy sequence call. MinION sequence read lengths range from several hundred bases to hundreds of thousands of bases, and are essentially limited by the DNA preparation type and delivery to the pore. Early versions of the MinION sequencing chemistry and base calling produced sequences with relatively high error rates (22-35%) when compared to both NGS (<2%) and PacBio TGS (10-15%). The high error rate combined with relatively low sequencing yields per flow cell (<1 Gb/flow cell) led some researchers to underestimate the potential for this technology. Recent advances in the MinION chemistry and base calling have greatly improved accuracy (5-10% error rate) and yield (2-5 Gb/flowcell) (J. Tyson unpublished results). To date, most studies have used the MinION to sequence small genomes or to partially survey larger genomes to assess chromosomal structure and copy number variations (Loman et al. 2015, Goodwin et al. 2015, Norris et al. 2016, Wei & Williams 2016,). The long read lengths of this technology, combined with recent improvements in performance make the MinION a viable option for whole genome sequencing of complex metazoan genomes.

The *Caenorhabditis elegans* genome was the first metazoan genome to be completely sequenced (C. elegans Sequencing Consortium 1998) and is an excellent model genome for assessing new whole genome sequencing technologies. The *C. elegans* genome is complex with many different types of local and dispersed repeat sequences in both intronic and intergenic regions and only 27% of the genome residing in exons. Local repeats range from short repetitive sequences such as homopolymeric G tracts (Zhao et al. 2007) to large tandem repeats spanning tens of kilobases. The most common dispersed repeats are derived from transposable elements and constitute approximately 12% of the genome (Bessereau 2006). *C. elegans* transposons range in size from 1-3 kb and can confound genomic assemblies as transposon sequences are larger than NGS and Sanger sequencing reads resulting in ambiguous mapping positions. The assembly of the *C. elegans* reference genome was made possible by the construction of a high quality physical map of cosmids and yeast artificial chromosomes (Coulson et al. 1988; Coulson et al. 1995; Coulson et al. 1991), which were individually shotgun sequenced and manually finished to bridge gaps and ambiguous regions to generate a complete high quality reference genome (C. elegans Sequencing Consortium 1998). The high-quality reference genome has facilitated studies using NGS re-sequencing to aid identifying new mutants and analysing mutational profiles (Thompson et al. 2013; Meier et al. 2014). Nevertheless, the potential for assembly discrepancies using NGS in sequence determination remain and NGS is not optimal for *de novo* genome assembly or understanding large structural alterations such as those associated with cancer and complex genetic disorders.

Here, we report the sequencing and *de novo* assembly of a 100 Mb *C. elegans* genome exclusively from MinION reads. For comparison, we used both MinION and Illumina platforms to sequence the genome of a *C. elegans* strain containing both a complex genome rearrangement generated by acetaldehyde mutagenesis and a ballistic-mediated insertional event of a GFP-containing plasmid (*him-9(e1487);ruIs32*). The *de novo* assembly of MinION data generated a remarkably complete genome in 145 contigs that cover >99% of the reference genome with an N50 contig length of 1.22 Mb. Furthermore, the sequence assembly elucidated the complex structure of the rearrangement and insertion events that were not readily apparent in the Illumina data. The strategy of comparison to Illumina data was however useful to improve the sequence accuracy of the MinION assembly. Together, this hybrid sequencing approach generated a physical map with high sequence accuracy and further represents the largest genome assembled exclusively from MinION data.

## Materials and Methods

### Nematode culture and DNA extraction

Nematodes were cultured as previously described (Brenner 1974). The *him-9(e1487)* II; *unc-119(ed3) ruIs32[pie-1p::GFP::H2B + unc-119(+)]* III strain was constructed by mating CB1487 *him-9(e1487)* males to AZ212 *unc-119*(*ed3*) *ruIs32* III hermaphrodites. F1 heterozygotes were selfed and *him-9; ruIs32* homozygotes isolated. Worms were grown to starvation for sequencing on NGM plates and harvested by washing with M9 buffer and pelleted in 15-ml centrifuge tubes. Buffer was removed by two washes with sterile distilled water, centrifugation and aspiration.

The worm pellet was resuspended in 300 μl of lysis buffer (200 mM NaCl, 100 mM Tris-HCl pH 8.5, 50 mM EDTA pH 8.0, 0.5% SDS, 0.1 mg/ml proteinase K) and frozen at −80 °C. Frozen pellets were incubated at 60 °C for 1-3 hours followed by 95°C for 20 minutes. RNase A (0.1 mg/ml) was added and lysed worms were incubated at 37°C for 1 hour. DNA was prepared by standard phenol/chloroform extraction and DNA was resuspended in 10 mM Tris pH 8.0.

### Library Preparation and MinION Sequencing

The Chip84-Chip90 MinION flowcells were run using libraries prepared with the SQK-NSK007 Nanopore Sequencing Kit R9 version. Chip94 & Chip95 libraries were prepared using the SQK-RAD001 Rapid Sequencing Kit I R9 version. The standard protocols from Oxford Nanopore Technologies were used with the following modifications. For SQK-NSK007 libraries, purification of DNA after the FFPE treatment step was done using 0.4x AMPureXP beads. Prior to adapter ligation each elution step off the AMPureXP beads was performed using 10 mM Tris pH 8.0, instead of water, at 37°C for 5 mins. The starting amounts of gDNA ranged between 0.8 μg and 2.0 μg (see Table S1 for details). Priming of individual flowcells with running buffer (2× 500 μl) and sequencing library top ups (150 μl) were performed at times detailed in Table 1. Flowcells were run for ∼48 hrs using custom device tuning scripts. The tuning scripts provide event yield monitoring aimed at maintaining data throughput through initiation of a maximal pore channel assignment/usage strategy and optimal bias-voltage selection via methods outlined below.

#### Modified MinION running scripts

After initial start of a MinION sequencer run using the standard ONT sequencing scripts, MinION sequencing control was shifted to a custom MinKNOW MinION script to enhance pore utilisation and increase data yields. This custom script adjusted a number of run/flowcell metrics and parameters including-

1) Initiation of a bias-voltage re-selection and active pore re-population into active channels when the hourly event yields falls below a threshold. This threshold was set at 67% of the first hour of each particular sequencing segment, and generally ran for 2-5 hours per segment.
2) Identifying the bias-voltages that provide the greatest number of active pores by scanning a voltage range (20-30 mV in increments of −10 mV) and using this for active pore channel reassignment. A newly selected bias-voltage acts as the starting point for subsequent scans, and provides an active pore “tracking” ability as the required bias-voltage magnitude increases during a run with the electrochemical gradient decay of individual wells. This re-assignment also provides access to the full 2048 possible wells repeatedly throughout a run.
3) Selecting a lower magnitude bias-voltage wherein the active pore numbers are within 10% of the peak voltage. Keeping greater pore numbers active using these approaches results in wells/pores running for different periods of time and the bias-voltage requirement to drive the same current through a pore increase and diverge with use. This is because the electrochemical gradient of active wells/pores decays at a greater rate than that of inactive wells/pores. By using off peak, lower magnitude bias-voltage selection, a measure of pore population containment or “shepherding” is provided by moving lower magnitude voltage requiring pores into the rest of the population.

For further details on these device running script modifications see, https://community.nanoporetech.com/posts/r9-tuning-scripts-for-mink, and links contained within.

### Base-calling MinION Sequencing Reads

All reads generated from the MinION sequencing device were base-called using the cloud-based Metrichor service provided by Oxford Nanopore. Details of specific versions can be found in Table S1 for the different runs. DNA sequences were extracted from individually called reads using simple python scripts and combined in a single fasta file format of a particular strand sequence type. Runs using the SQK-NSK007 (2D) kit generated template and a fraction of complement and 2D sequence from individual reads. SQK-RAD001 (1D) Library runs generated template (1D) reads. Filtering based on a quality metric by Metrichor divided the reads additionally into ‘pass’ and ‘fail’ categories. Some or all these sequence containing files for each run were then used for genome assembly as indicated.

## Genome assembly and evaluation

### SPAdes Illumina assembly

For Illumina data, low quality bases were trimmed from both ends using seqtk (lh. lh3/seqtk. *GitHub* Available at: https://github.com/lh3/seqtk. (Accessed: 26th November 2016)). The threshold used for trimming was a quality score of 30. We performed an Illumina only assembly using SPAdes (Nurk et al. 2013) in its default settings with kmer sizes 21, 51, and 71. The number of threads used was 32 with a memory of 156 GB.

### Canu nanopore assembly

We pooled all the nanopore data (pass and fail; 1D and 2D) and filtered out reads below a size of 1 kb to avoid overlap detection issues. The remainder of reads (∼6.03 gigabases) were assembled using Canu (Koren et al. 2016) in its default settings.

### Assembly correction

The Canu assemblies were corrected using Pilon (Walker et al. 2014) using recommended settings to polish for variants and homopolymers. We did not perform quality filtering on the Illumina data and all of the ∼8.04 gigabases of short read sequence was used.

### Nanopore data and assembly evaluation

Nanopore reads and assemblies were evaluated by alignment against the reference worm genome using marginAlign version 0.1 (with BWA-MEM (Li 2013)) (Jain et al. 2015). Alignment and error statistics were computed using marginStats version 0.1 (Jain et al. 2015). To better estimate the errors for nanopore data, we improved the alignments using marginAlign EM (Jain et al. 2015). We also performed secondary evaluations using LASTZ alignments to align draft assemblies against the reference.

We evaluated the assembly quality by using QUAST (Gurevich et al. 2013) with recommended settings and the reference worm genome. We compared changes in indels and mismatches for the Canu assemblies before and after Pilon correction. We evaluated the assemblies based on: 1) the total number of aligned bases in the assembly; 2) the number of mismatches; and 3) the total number of bases contained in indels.

## Results

### Sequencing a *C. elegans* strain with complex rearrangements

To assess the feasibility of using MinION sequencing to both generate *de novo* whole genome assemblies of large genomes and delineate complex rearrangements, we constructed for sequencing a *C. elegans* strain, *him-9(e1487)* II; *unc-119(ed3) ruIs32[pie-1p::GFP::H2B + unc-119(+)]* III, that contained two homozygous complex rearrangements. The *him-9(e1487)* mutation was induced by acetaldehyde mutagenesis (Hodgkin et al. 1979) and is a complex duplication and insertion event that disrupts the predicted *C. elegans XPF* orthologue *xpf-1* (Youds et al. 2006). Previous analysis of *him-9(e1487)* determined that the insertion contained duplicated sequence from the *mab-3* gene (N.J. O’Neil, unpublished data). However, the complex nature of the mutation stymied attempts to determine the exact structure of the rearrangement. The *ruIs32* insertion is a low-copy number insertion that was generated by biolistic transformation of the plasmid pAZ132 [*Ppie-1::GFP::H2B::pie-1*] (derived from pJH4.52) and a plasmid containing *unc-119* [*unc-119(+)*] into an *unc-119(ed3)* mutant (Wormbase). Genomic DNA was prepared from the *him-9(e1487)* II; *ruIs32* III strain and sequenced by both a MinION sequencer using R9 chemistry and Illumina sequencing using 300 base paired-end reads.

Sequencing runs from six MinION flow cells using the R9.0 (4x) and R9.3 (2x) pore flowcell types were produced on the MinION device controlled by custom tuning scripts (see Methods). This resulted in 1.1M individual reads up to 123,159 bases in length (mean = 4,801) and containing 5.33Gb of 1D bases. An additional 1Gb of 2D sequence was generated from the paired template and complement 1D reads produced from the R9.0 flow cells using the SQK-NSK007 2D chemistry and Metrichor basecaller. Details of the individual MinION sequencing runs and the MinION sequencing chemistries used can be found in (Figure 1) and (Table 1). Significant improvements in the 1D sequence quality of individual reads was observed when comparing R9.3 to R9.0 (Figure S1). This evolution on a path to the present day R9.4 pore provided significant advances in the simpler production of higher quality 1D reads approaching the percentage read identity of the previous generation 2D reads. Nanopore read quality was measured by alignment to the *C. elegans* reference genome. The median identity for pass 1D reads from the 1D runs that used SQK-RAD001 Rapid Sequencing Kit I R9 version was ∼93%. The median identities for pass 2D reads from the 2D runs that used SQK-NSK007 Nanopore Sequencing Kit R9 version (Chips 84-90) ranged between 90-95%. Illumina sequencing produced 14,041,499 paired end reads totaling ∼8.04 Gb of sequence (Table 2). Alignment of the MinION sequence reads to the *C. elegans* reference genome demonstrated that most of the genome was well covered (∼50X coverage) and identified an apparent ∼2 Mb duplication on chromosome III (Figure 2).

**Figure 1.**
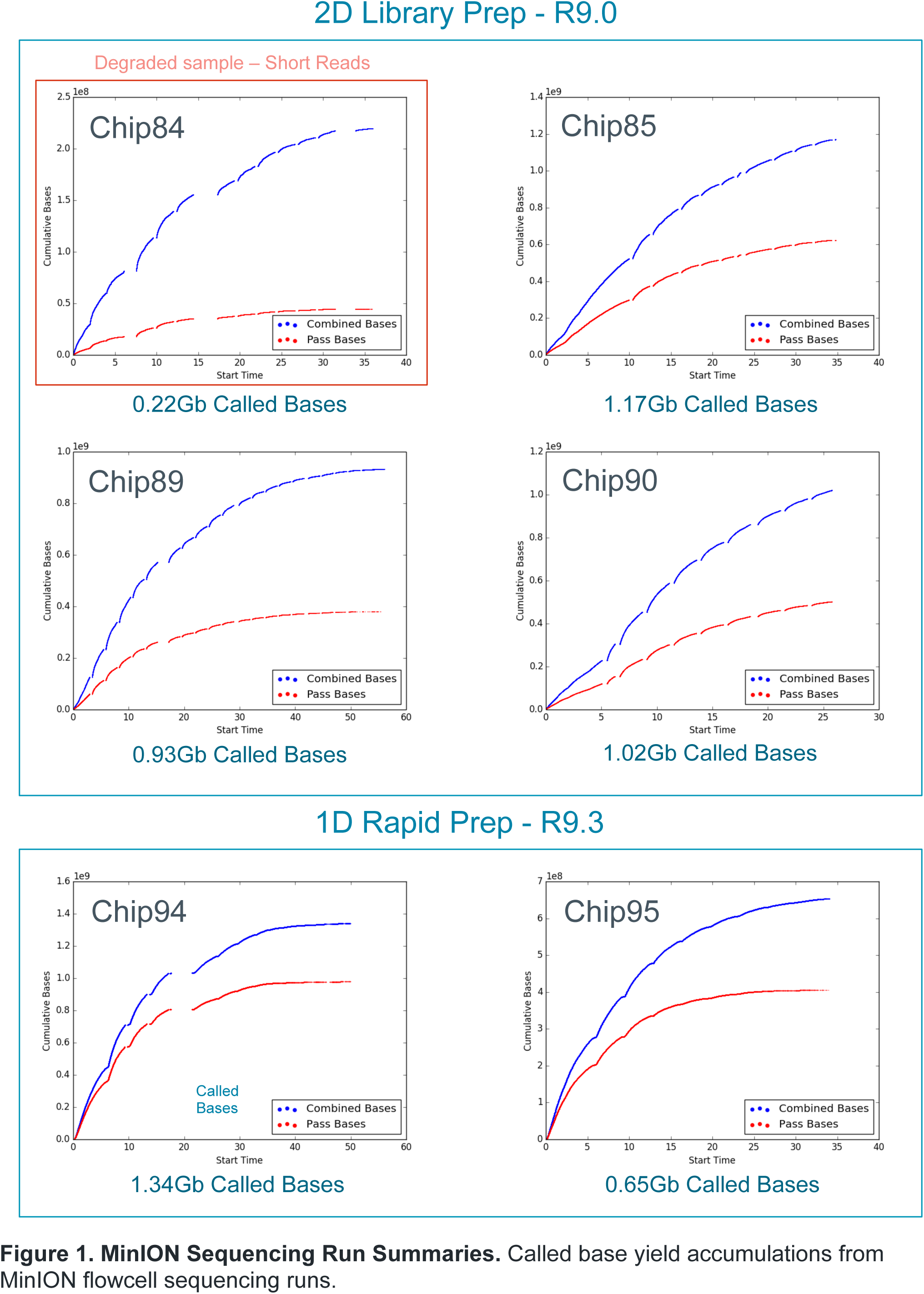
MinION Sequencing Run Summaries. Called base yield accumulations from MinION flowcell sequencing runs. Dot plots of LASTZ alignments on MinION Pilon polished contigs to the *C. elegans* reference chromosomes. * denotes contig that aligns to chromosome II and V.

**Table 1.** MinION Sequencing Data Summary. Individual library preparation and run statistics for MinION sequencing of *C.elegans him-9* mutant.

|  | Chip84 | Chip85 | Chip89 | Chip90 | Chip94 | Chip95 |
| --- | --- | --- | --- | --- | --- | --- |
| Library Kit | SQK-NSK007 | SQK-NSK007 | SQK-NSK007 | SQK-NSK007 | SQK-RAD001 | SQK-RAD001 |
| Flowcell ID | FAD11292 | FAD11271 | FAD11308 | FAB23558 | FAD24831 | FAD24186 |
| Pore Type | R9.0 | R9.0 | R9.0 | R9.0 | R9.3 | R9.3 |
| Genomic DNA Start Material | 749ng | 1650ng | 1978ng | 1978ng | 864ng into 20ul | 1152ng into 20ul |
| Volume Loaded | 12ul 0hrs, 12ul 0.5hrs | 12ul 0hrs, | 12ul 0hrs | 6ul 0hrs, 6ul 24hrs | 11ul 0hrs, 11ul 24hrs | 11ul 0hrs, 11ul 24hrs |
| Preseq Mix Conc. | 7.14ng/ul | 5.34ng/ul | 8.56ng/ul | 8.56ng/ul | na | na |
| PlatQC Pores | 1155//486,397,217,55 | 1036//467,357,170,42 | 1060//428,340,222,70 | 1404//489,444,330,141 | 1277//494,423,271,89 | 1361//504,448,312,97 |
| **Total Reads** | **198,913** | **184,600** | **142,374** | **155,508** | **268,004** | **161,378** |
| Pass Reads | 36,949 | 68,204 | 34,816 | 54,892 | 180,938 | 89,376 |
| Av Template Read Length | 831 | 4,052 | 4,486 | 4,430 | 5,001 | 4,049 |
| Av Complement Read Length | 272 | 2,281 | 2,059 | 2,125 | na | na |
| Av 2D Read Length | 171 | 2,207 | 1,826 | 1,972 | na | na |
| Max Pass Template Read size | 9,460 | 29,221 | 44,484 | 32,910 | 108,915 | 123,159 |
| Max Pass Complement Read size | 9,130 | 28,814 | 34,881 | 31,458 | na | na |
| Max Pass 2D Read size | 9,740 | 29,681 | 44,405 | 33,553 | na | na |
| Top 25% Templates > | 904 | 6,106 | 6,858 | 6,622 | 6,390 | 4,717 |
| Top 25% Complements > | 305 | 4,233 | 2,721 | 2,896 | na | na |
| Top 25% 2D > | 273 | 4,454 | 1,987 | 2,533 | na | na |
| **Total Called bases** | **219,411,804** | **1,169,061,304** | **931,927,653** | **1,019,278,312** | **1,340,369,063** | **653,447,905** |
| 2D Pass Bases | 22,208,070 | 323,598,976 | 195,759,566 | 258,533,514 | na | na |
| 2D fail Bases | 11,877,828 | 83,731,694 | 64,225,734 | 48,089,637 | na | na |
| 2D Bases | 34,085,898 | 407,330,670 | 259,985,300 | 306,623,151 | na | na |
| Base Calling | 2D Chimera 1.22.10 | 2D Chimera 1.22.10 | 2D Chimera 1.22.10 | 2D Chimera 1.22.10 | 1D Chimera 1.22.10 | 1D Chimera 1.22.10 |
| MinkNOW version | 0.51.3.55 | 0.51.3.55 | 0.51.3.55 | 0.51.3.55 | 1.0.5 | 1.0.5 |
| Comment | Small Frags degraded Library |  |  | SpotON Flowcell |  |  |

**Figure S1.**
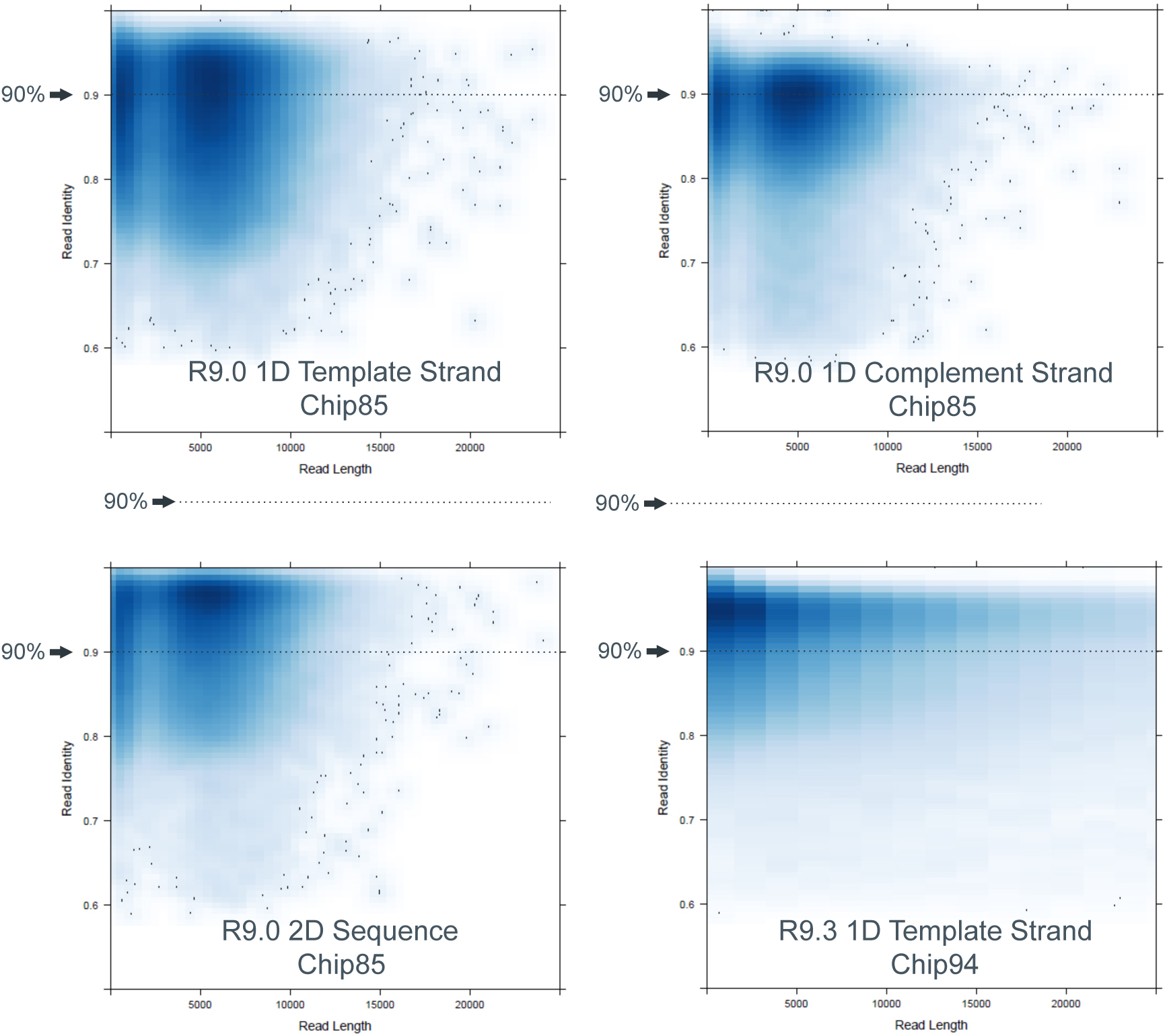
MinION Individual Read Accuracies vs Read Length. Example plots for both R9.0 2D and R9.3 1D chemistries. Individual pass read accuracies from aligned regions calculated using bwa alignment to the reference sequence and plotted against individual read length.

**Figure 2.**
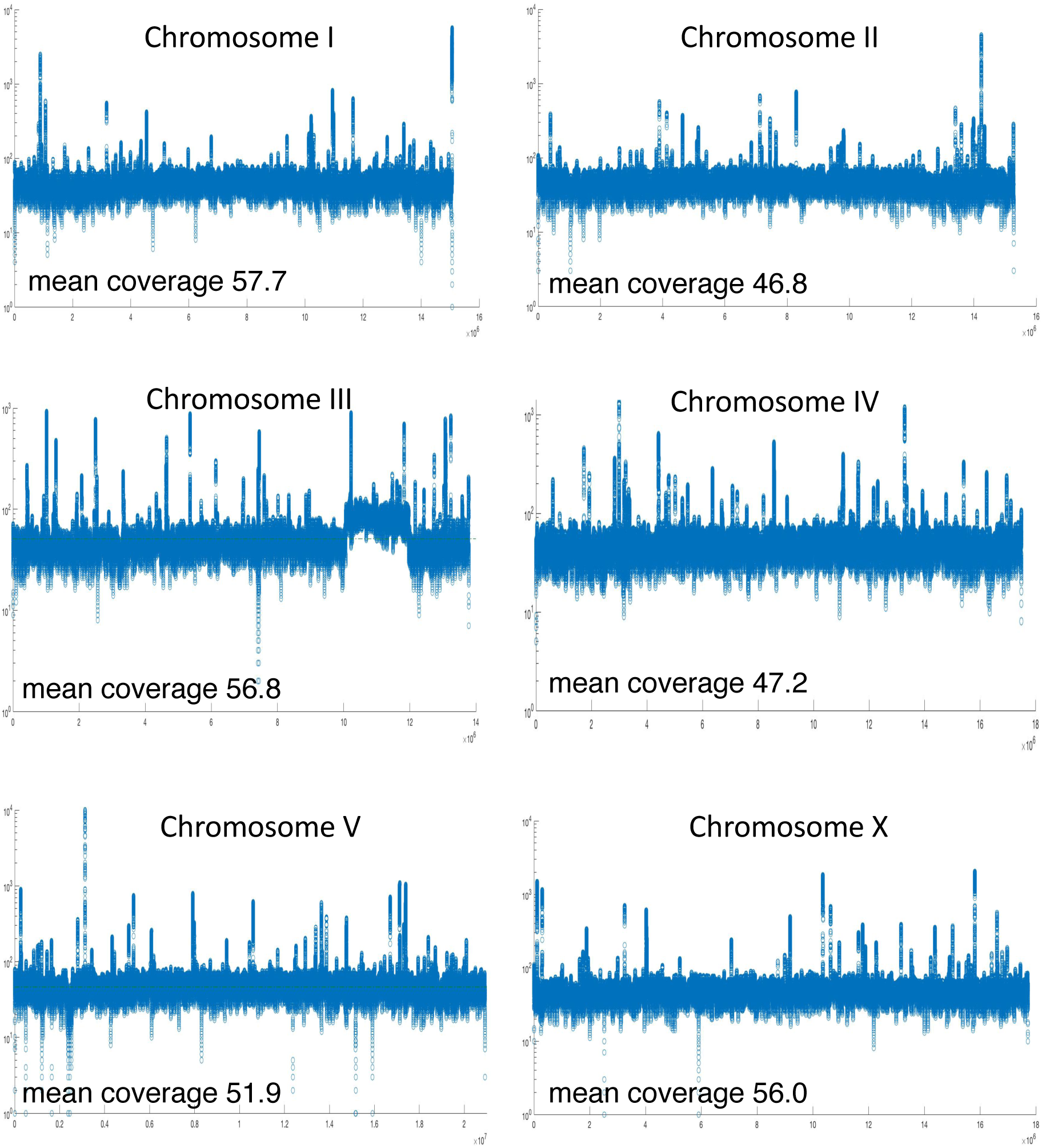
MinION read coverage (log scale) aligned to reference genome. Note the duplication between 10,062,096 and 11,973,739 on chromosome III.

**Table 2.**
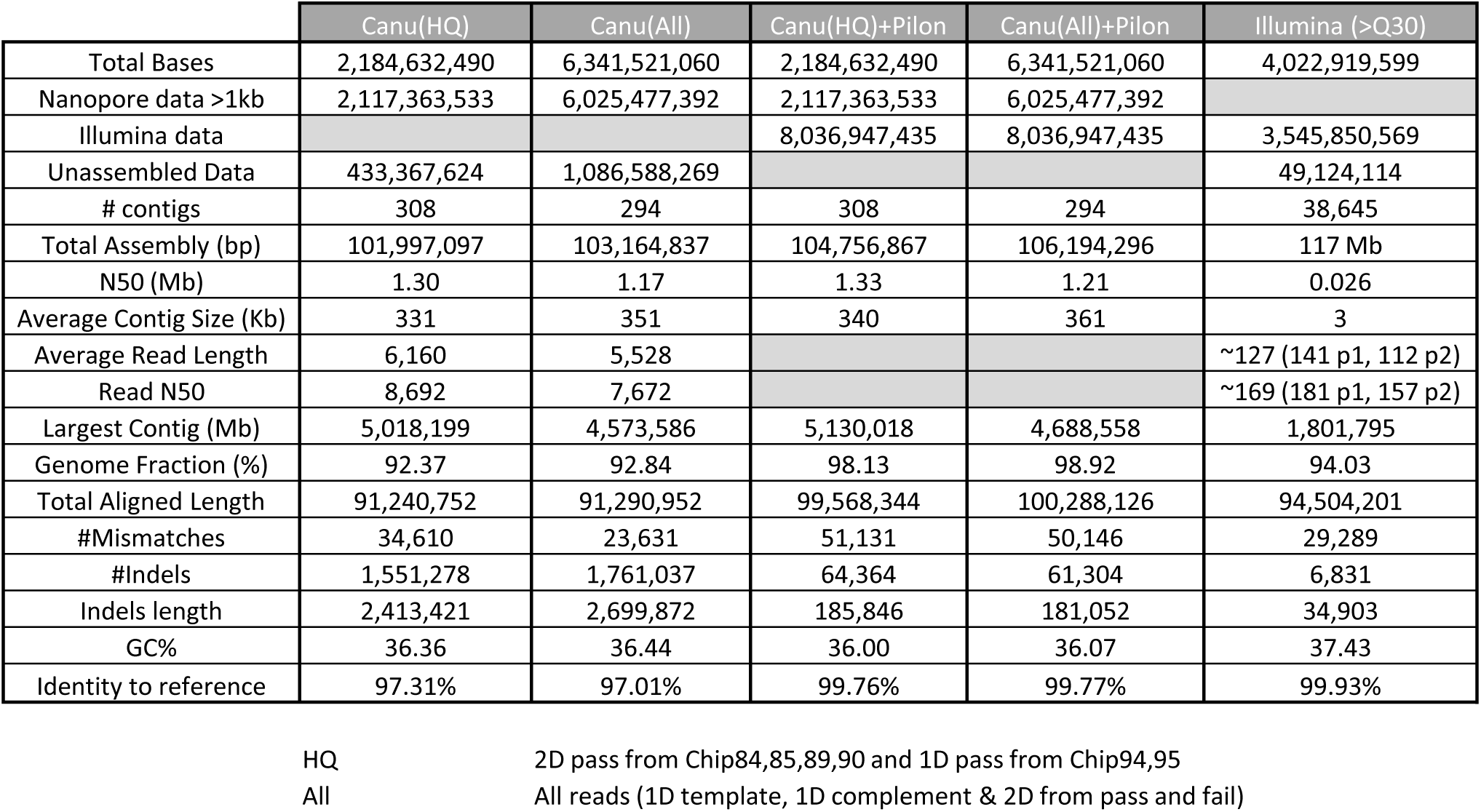
Illumina sequencing Read Data

### *de novo* assembly of the *C. elegans* genome from MinION and Illumina sequence reads

1D and 2D sequence reads were filtered by size to exclude reads < 1kb and Canu (Korin et al 2016) a genome assembler designed for high-noise single-molecule sequencing was used to assemble contigs. A Canu assembly was performed using all “pass” and “fail” reads from all prepared library types to generate an assembly containing 294 contigs ranging in length from 3,695 to 4,573,586 bases with an average contig size of 351 kb. The Canu contigs were polished using Pilon and the Illumina sequence data (Table 3). 145 of the contigs containing 101,982,548 bases had significant homology to *C. elegans* (Table 4). Most of the non-*C. elegans* contigs were homologous to bacterial genomes consistent with the bacteria present on the NGM petri plates used to grow the nematodes. The mean length for *C. elegans* matching contigs was 703,328 bases compared to a mean contig length of 28,266 bases for the 149 non-*C. elegans* contigs, likely due to lower coverage. The 145 *C. elegans* contigs were aligned to the WBcel235 release of the *C. elegans* reference genome using LASTZ aligner (Harris, R.S. 2007). The alignment of contigs to the six chromosomes and the *C. elegans* mitochondrial genome covered >99% of the reference genome with >97% identical sites (Table 5; Figure 3). There were no large gaps in coverage of the reference genome. Three contigs, aligned to two different chromosomes resulting in apparent discontinuities. These hybrid contigs were most likely misassembled contigs as no other evidence suggests that these regions have been translocated.

**Figure 3.**
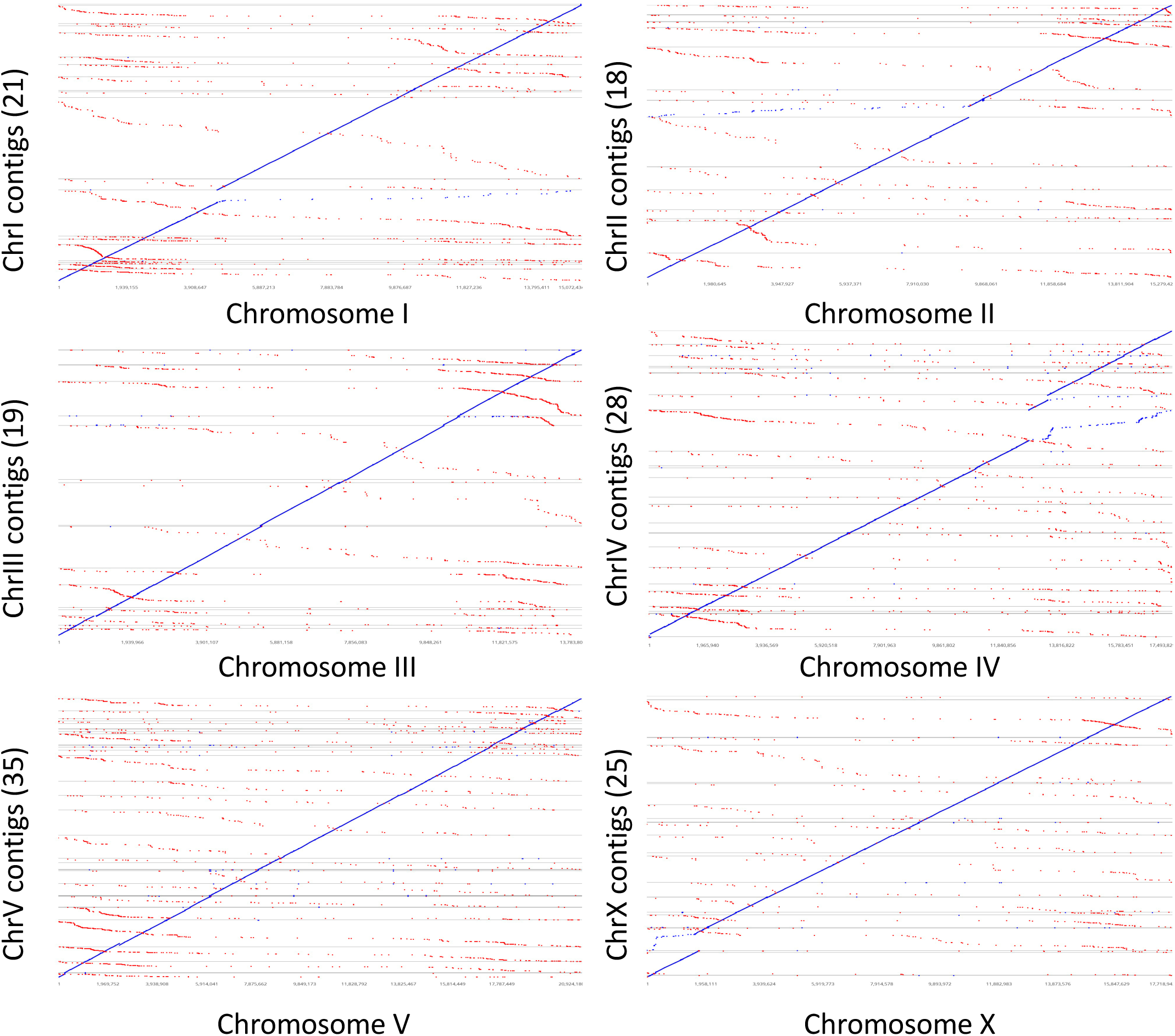
LASTZ alignments of MinION contigs to *C. elegans* reference chromosomes. Discontinuities are due to contigs mapping to two different chromosomes. We have included these hybrid contigs on both chromosomes to allow for alignment. These hybrid contigs are most likely the result of contig assembly errors in highly repetitive regions.

**Table 3.**
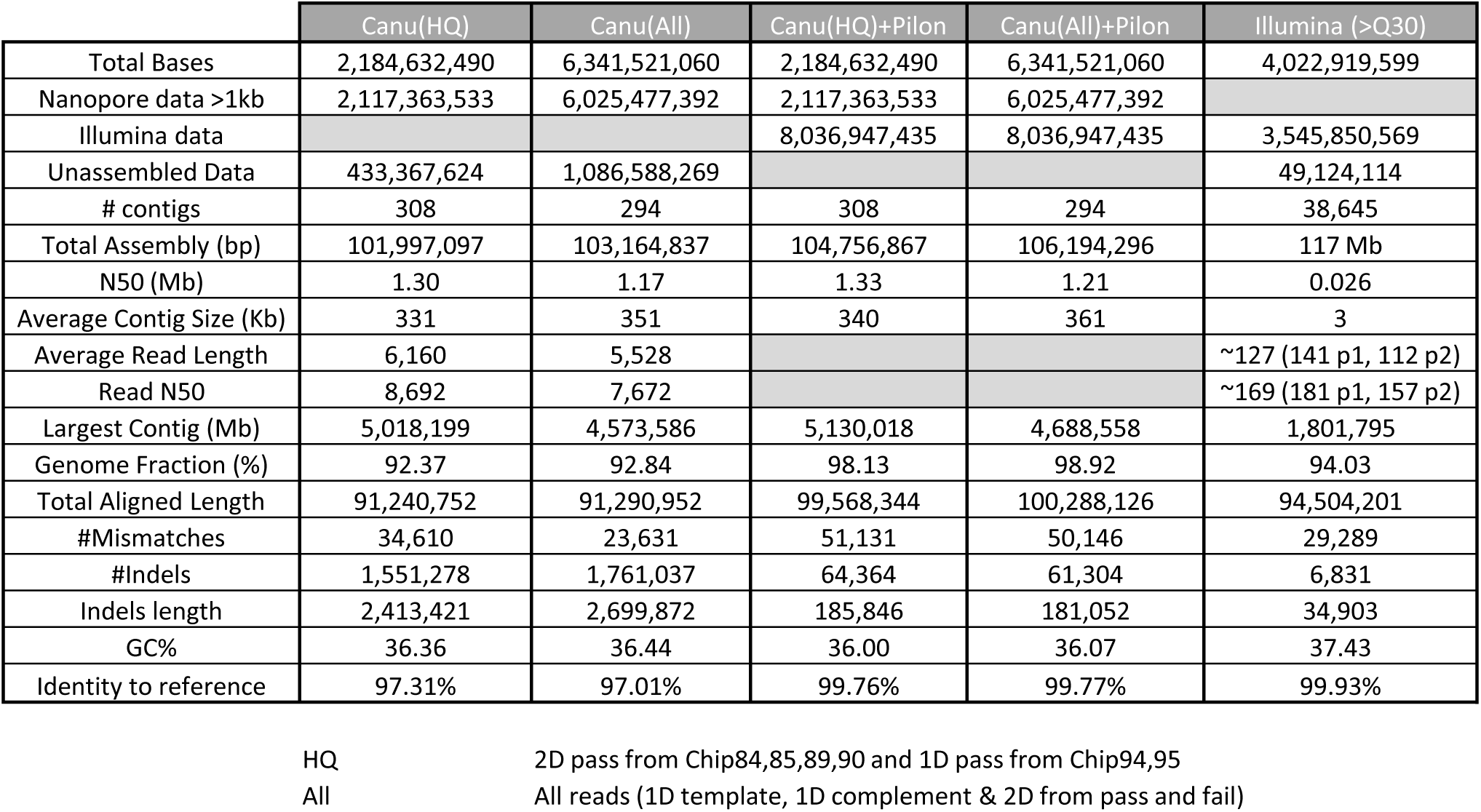
*C.elegans him-9* mutant Genome Assemblies and Polishing. Different genome assemblies using Nanopore, Illumina and combined data sets. (HQ) 2D pass from Chip84, 85, 89, 90 and 1D pass from Chip94, 85. (All) All reads, 1D template, 1D complement & 2D from pass and fail catagories.

**Table 4.** Worm Contigs

| Chromosome I |  |  | Chromosome II |  |  | Chromosome III |  |  | Chromosome IV |  |  | Chromosome V |  |  | Chromosome X |  |  | Mitochondrial chromosome |  |  |
| --- | --- | --- | --- | --- | --- | --- | --- | --- | --- | --- | --- | --- | --- | --- | --- | --- | --- | --- | --- | --- |
| CONTIG | LENGTH | PILON LENGTH | CONTIG | LENGTH | PILON LENGTH | CONTIG | LENGTH | PILON LENGTH | CONTIG | LENGTH | PILON LENGTH | CONTIG | LENGTH | PILON LENGTH | CONTIG | LENGTH | PILON LENGTH | CONTIG | LENGTH | PILON LENGTH |
| 0128 | 641,577 | 667,188 | 0093 | 1,404,993 | 1,452,753 | 1829 | 322,164 | 335,057 | 0069 | 57,949 | 59,969 | 1815 | 289,924 | 300,768 | 0291 | 105,703 | 110,238 | 0111 | 26,495 | 26,994 |
| 0162 | 249,830 | 261,633 | 0066 | 1,825,969 | 1,884,875 | 1827 | 145,979 | 151,731 | 0064 | 1,470,161 | 1,523,240 | 1816 | 47,237 | 47,480 | 1808 | 968,329 | 995,374 |  |  |  |
| 0146 | 101,839 | 105,941 | 0190 | 145,554 | 149,288 | 0157 | 449,989 | 464,710 | 1775 | 23,228 | 23,545 | 1817 | 827,816 | 855,414 | 1839 | 3,895 | 3,979 |  |  |  |
| 0155 | 90,893 | 94,925 | 0113 | 522,805 | 538,113 | 0044 | 286,814 | 299,041 | 1774 | 152,421 | 158,647 | 1793 | 1,083,739 | 1,121,912 | 1791 | 1,525,866 | 1,573,864 |  |  |  |
| 0060 | 910,770 | 955,944 | 1732 | 1,147,314 | 1,180,765 | 1753 | 106,886 | 110,089 | 0022 | 309,830 | 325,753 | 0039 | 1,999,640 | 2,060,681 | 0056 | 118,963 | 126,367 |  |  |  |
| 0144 | 285,340 | 305741 | 1776 | 1,333,788 | 1,365,313 | 0063 | 1,054,603 | 1,105,484 | 0084 | 936,097 | 976,525 | 1802 | 937,727 | 968,247 | 0089 | 48,192 | 49,234 |  |  |  |
| 0041 | 147,174 | 159,766 | 1777 | 15,658 | 15,848 | 0121 | 780,480 | 817,746 | 0019 | 475,203 | 505,301 | 1803 | 34,408 | 34,549 | 1759 | 836,347 | 862,726 |  |  |  |
| 1812 | 597,398 | 614,699 | 1778 | 2,891,883 | 2,957,296 | 0031 | 1,971,186 | 2,029,091 | 1787 | 1,055,217 | 1,091,923 | 1742 | 793,648 | 813,478 | 1757 | 174,755 | 180,694 |  |  |  |
| 1811 | 19,219 | 19,306 | 1789 | 15,231 | 15,270 | 1735 | 55,591 | 55,699 | 0029 | 1,325,883 | 1,363,306 | 1765 | 18,063 | 18,253 | 1730 | 1,014,182 | 1,041,746 |  |  |  |
| 1725 | 4,573,586 | 4,688,558 | 0023 | 595,982 | 609,952 | 1736 | 2,020,242 | 2,069,914 | 1750 | 889,415 | 913,229 | 1807 | 27,100 | 27,102 | 0456 | 11,254 | 11,286 |  |  |  |
| 1729 | 318,637 | 327,171 | 1766 | 2,478,930 | 2,559,627 | 1798 | 159,446 | 162,596 | 1751 | 39,749 | 40,436 | 1835 | 22,957 | 23,062 | 0075 | 1,766,377 | 1,813,766 |  |  |  |
| 1727 | 83,750 | 85,600 | 0009 | 1,108,459 | 1,153,611 | 1783 | 2,577,988 | 2,640,414 | 0042 | 627,979 | 642,707 | 1764 | 873,000 | 893,363 | 0211 | 205,924 | 211,400 |  |  |  |
| 0018 | 750,497 | 770,051 | 0119 | 327,432 | 341,061 | 0040 | 447,466 | 460,463 | 1796 | 1,195,712 | 1,223,010 | 1762 | 967,542 | 989,686 | 0102 | 1,178,956 | 1,207,894 |  |  |  |
| 0136 | 681,659 | 708,475 | 1819 | 18,180 | 18,661 | 0065 | 1,618,827 | 1,687,451 | 1795 | 18,456 | 18,575 | 1810 | 80,862 | 82,488 | 1780 | 848,498 | 870,998 |  |  |  |
| 0152 | 422,339 | 438,644 | 1806 | 371,509 | 383,023 | 1790 | 771,310 | 804,548 | 1794 | 460,691 | 471,631 | 1761 | 19,250 | 19,316 | 1779 | 23,740 | 24,063 |  |  |  |
| 0097 | 1,353,294 | 1,397,343 | 1805 | 24,717 | 25,461 | 1836 | 17,002 | 17,100 | 0099 | 1,275,584 | 1,304,238 | 0092 | 505,374 | 516,537 | 1744 | 259,389 | 265,556 |  |  |  |
| 0156 | 394,056 | 410,474 | 1804 | 550,140 | 573,165 | 1788 | 692,834 | 717,085 | 0124 | 649,249 | 663,972 | 1821 | 328,510 | 335,880 | 0114 | 898,142 | 919,234 |  |  |  |
| 0068 | 110,325 | 116,857 |  |  |  | 1758 | 3,917 | 3,923 | 1740 | 102,549 | 104,785 | 1737 | 1,751,778 | 1,788,507 | 0074 | 1,406,913 | 1,439,548 |  |  |  |
| 1813 | 1,030,425 | 1,069,619 |  |  |  | 0808 | 7,998 | 8,057 | 0011 | 944,601 | 967,191 | 1739 | 1,874,708 | 1,915,981 | 1734 | 111,328 | 114,321 |  |  |  |
| 1814 | 75,058 | 75,451 |  |  |  |  |  |  | 0003 | 2,682,200 | 2,772,593 | 0107 | 1,110,630 | 1,136,009 | 1767 | 2,448,979 | 2,515,054 |  |  |  |
|  |  |  |  |  |  |  |  |  | 1771 | 1,395,592 | 1,441,393 | 0095 | 1,073,058 | 1,099,497 | 1769 | 526,550 | 540,649 |  |  |  |
|  |  |  |  |  |  |  |  |  | 0055 | 972,204 | 999,639 | 1801 | 1,916,623 | 1,966,771 | 1745 | 23,913 | 24,056 |  |  |  |
|  |  |  |  |  |  |  |  |  | 1770 | 24,088 | 24,306 | 0015 | 381,581 | 391,181 | 1746 | 2,477,698 | 2,555,070 |  |  |  |
|  |  |  |  |  |  |  |  |  | 1773 | 298,435 | 308,756 | 1826 | 249,592 | 254,471 | 1747 | 6,362 | 6,364 |  |  |  |
|  |  |  |  |  |  |  |  |  | 0161 | 105,733 | 109,044 | 1824 | 136,416 | 138,132 | 1748 | 244,817 | 252,069 |  |  |  |
|  |  |  |  |  |  |  |  |  | 1760 | 694,777 | 726,700 | 1763 | 19,006 | 19,356 |  |  |  |  |  |  |
|  |  |  |  |  |  |  |  |  | 0081 | 710,949 | 738,400 | 1785 | 813,298 | 843,246 |  |  |  |  |  |  |
|  |  |  |  |  |  |  |  |  | 0123 | 877,321 | 907,738 | 1756 | 173,877 | 179,974 |  |  |  |  |  |  |
|  |  |  |  |  |  |  |  |  |  |  |  | 0017 | 171,054 | 177,803 |  |  |  |  |  |  |
|  |  |  |  |  |  |  |  |  |  |  |  | 0147 | 64,693 | 67,991 |  |  |  |  |  |  |
|  |  |  |  |  |  |  |  |  |  |  |  | 1823 | 350,901 | 365,730 |  |  |  |  |  |  |
|  |  |  |  |  |  |  |  |  |  |  |  | 0165 | 208,425 | 218,028 |  |  |  |  |  |  |
|  |  |  |  |  |  |  |  |  |  |  |  | 0175 | 167,421 | 172,916 |  |  |  |  |  |  |
|  |  |  |  |  |  |  |  |  |  |  |  | 0142 | 571,657 | 596,057 |  |  |  |  |  |  |
|  |  |  |  |  |  |  |  |  |  |  |  | 0117 | 922,679 | 955,919 |  |  |  |  |  |  |
| Total CONTIG | 12837666 | 13273386 |  | 14,778,544 | 15,224,082 |  | 13,490,722 | 13,940,199 |  | 19,771,273 | 20,406,552 |  | 20,814,194 | 21,395,785 |  | 17,235,07 | 17,715,550 |  | 26,495 | 26,994 |
| Reference bp |  | 15,072,434 |  |  | 15,279,421 |  |  | 13,783,801 |  |  | 17,493,829 |  |  | 20,924,180 |  |  | 17,718,942 |  |  | 13,794 |

**Table 5.** LASTZ alignment of contigs to the Six chromosomes and the *C.elegans* mitochondrial genome.

| **Chromosome**<br>**(Size in bp)** | **Contigs after Pilon** |  |  |
| --- | --- | --- | --- |
|  | Pairwise<br>Identity (%) | Identical<br>Sites (%) | Reference<br>Coverage (%) |
| I (15,072,434) | 80.9 | 95.0 | 99.8 |
| II (15,279,421) | 83.3 | 96.1 | 99.9 |
| III (13,783,801) | 84.5 | 94.4 | 99.2 |
| IV (17,493,829) | 85.3 | 96.3 | 99.8 |
| V (20,924,180) | 85.7 | 96.0 | 99.2 |
| X (17,718,942) | 84.1 | 97.5 | 99.5 |
| Mito (13,794) | 98.5 | 98.8 | 100 |

### MinION generated contigs unambiguously assigned transposon locations

To assess the quality of the assembly with respect to repetitive elements, we compared the number and position of Tc1 transposons in the *him-9; ruIs32* MinION generated contigs to the reference genome. Tc1 is a 1.6 kb transposon. NGS paired end reads are not sufficiently long to span the Tc1 transposons and cannot be unambiguously mapped resulting in a mapping quality score of 0 when aligned by the BWA aligner. Transposon number and position in the assembled contigs can be used as a measure of how effective MinION long reads are for spanning dispersed repeat regions. BLAST was used to align Tc1 reference sequences to the 145 worm contigs and the *C. elegans* reference genome. All described Tc1 elements were present in the MinION contigs and corresponded to their position in the reference genome based on the LASTZ alignments. (Table 6)

**Table 6.** Tc1 transposon numbers, locations and match compared to reference Genome.

| MinION Contigs |  |  |  |  | Reference Genome |  |  |  |
| --- | --- | --- | --- | --- | --- | --- | --- | --- |
| Contig Name | Chr | Length | % Pairwise Identity | % Query Coverage | Chr | Length | % Pairwise Identity | % Query Coverage |
| tig00000076 | I | 1,610 | 99.6 | 100 | I | 1,611 | 99.6 | 100 |
| tig00000097 | I | 852 | 99.3 | 53 | I | 852 | 99.3 | 53 |
| tig00000097 | I | 741 | 99.7 | 46 | I | 741 | 99.7 | 46 |
| tig00000148 | I | 1,532 | 98.5 | 100 | I | 1,611 | 99.6 | 100 |
| tig00000066 | II | 1,598 | 98.8 | 100 | II | 1,612 | 99.8 | 100 |
| tig00000093 | II | 1,598 | 98.9 | 100 | II | 1,611 | 99.9 | 100 |
| tig00000093 | II | 1,608 | 99.7 | 100 | II | 1,612 | 99.6 | 100 |
| tig00001732 | II | 862 | 99.7 | 53 | II | 861 | 99.7 | 53 |
| tig00001766 | II | 1,600 | 98.8 | 100 | II | 1,611 | 99.6 | 100 |
| tig00001766 | II | 1,605 | 99.2 | 100 | II | 1,611 | 99.6 | 100 |
| tig00001778 | II | 1,605 | 99.3 | 100 | II | 1,611 | 99.8 | 100 |
| tig00001778 | II | 1,609 | 99.1 | 100 | II | 1,611 | 99.4 | 100 |
| tig00000009 | II | 1,473 | 99.1 | 91 | II | 1,472 | 99.3 | 91 |
| tig00000063 | III | 1,613 | 99.4 | 100 | III | 1,611 | 99.6 | 100 |
| tig00001790 | III | 1,599 | 99.1 | 100 | III | 1,611 | 99.7 | 100 |
| tig00000055 | IV | 1,610 | 99.6 | 100 | IV | 1,610 | 99.6 | 100 |
| tig00000084 | IV | 1,582 | 98.0 | 100 | IV | 1,611 | 99.6 | 100 |
| tig00000099 | IV | 1,608 | 99.4 | 100 | IV | 1,611 | 99.6 | 100 |
| tig00001760 | IV | 1,602 | 99.2 | 100 | IV | 1,611 | 99.7 | 100 |
| tig00001793 | V | 1,511 | 99.0 | 94 | V | 1,522 | 99.7 | 94 |
| tig00001793 | V | 1,603 | 99.2 | 100 | V | 1,611 | 99.7 | 100 |
| tig00001785 | V | 1,595 | 98.6 | 100 | V | 1,611 | 99.4 | 100 |
| tig00000017 | V | 1,609 | 99.6 | 100 | V | 1,612 | 99.3 | 100 |
| tig00001801 | V | 1,607 | 99.3 | 100 | V | 1,611 | 99.6 | 100 |
| tig00001737 | V | 725 | 98.9 | 45 | V | 729 | 99.5 | 45 |
| tig00001737 | V | 1,610 | 99.1 | 100 | V | 1,610 | 99.6 | 100 |
| tig00000005 | V | 1,504 | 99.3 | 94 | V | 1,508 | 99.6 | 94 |
| tig00000039 | V | 1,602 | 99.1 | 100 | V | 1,611 | 99.8 | 100 |
| tig00000039 | V | 1,595 | 98.8 | 100 | V | 1,611 | 99.8 | 100 |
| tig00000039 | V | 1,602 | 99.3 | 100 | V | 1,611 | 99.6 | 100 |
| tig00001739 | V | 1,596 | 98.8 | 100 | V | 1,611 | 99.6 | 100 |
| tig00000211 | X | 1,602 | 99.1 | 100 | X | 1,611 | 99.6 | 100 |
| tig00000074 | X | 1,613 | 99.3 | 100 | X | 1,611 | 99.4 | 100 |

### MinION reads elucidated the structure of complex repetitive chromosome rearrangements

In addition to *de novo* genome assembly, long sequencing reads can also facilitate the delineation of complex chromosomal rearrangements that contain multiple breakpoints, duplications, deletions and repeated sequences. The *C. elegans* strain sequenced here possessed two different complex genome rearrangements: *him-9(e1487)* is an acetaldehyde-induced duplication-insertion event on chromosome II and *ruIs32* is a biolistic-mediated transgenic insertion on chromosome III. Previous data from oligo array hybridization, reverse transcriptase PCR, and inverse-PCR experiments suggested that *him-9(e1487)* was an insertion of approximately 20 kb of sequence from the *mab-3* region into the *xpf-1* gene. Assembly of MinION reads revealed a significantly more complex genomic rearrangement. While the insertion breakpoints were consistent with previous data, the insertion was larger than previously anticipated. In addition to the *mab-3* duplicated region, the insertion also included an inverted repeat of part of the *mab-3* duplication and the second exon of *xpf-1,* resulting in a much larger, more complex insertion than expected (Figure 4A). We were able to use paired end reads and copy number variations in the Illumina data to confirm the multiple breakpoints and copy number variations of the *mab-3* region (Figure 4B).

**Figure 4A.**
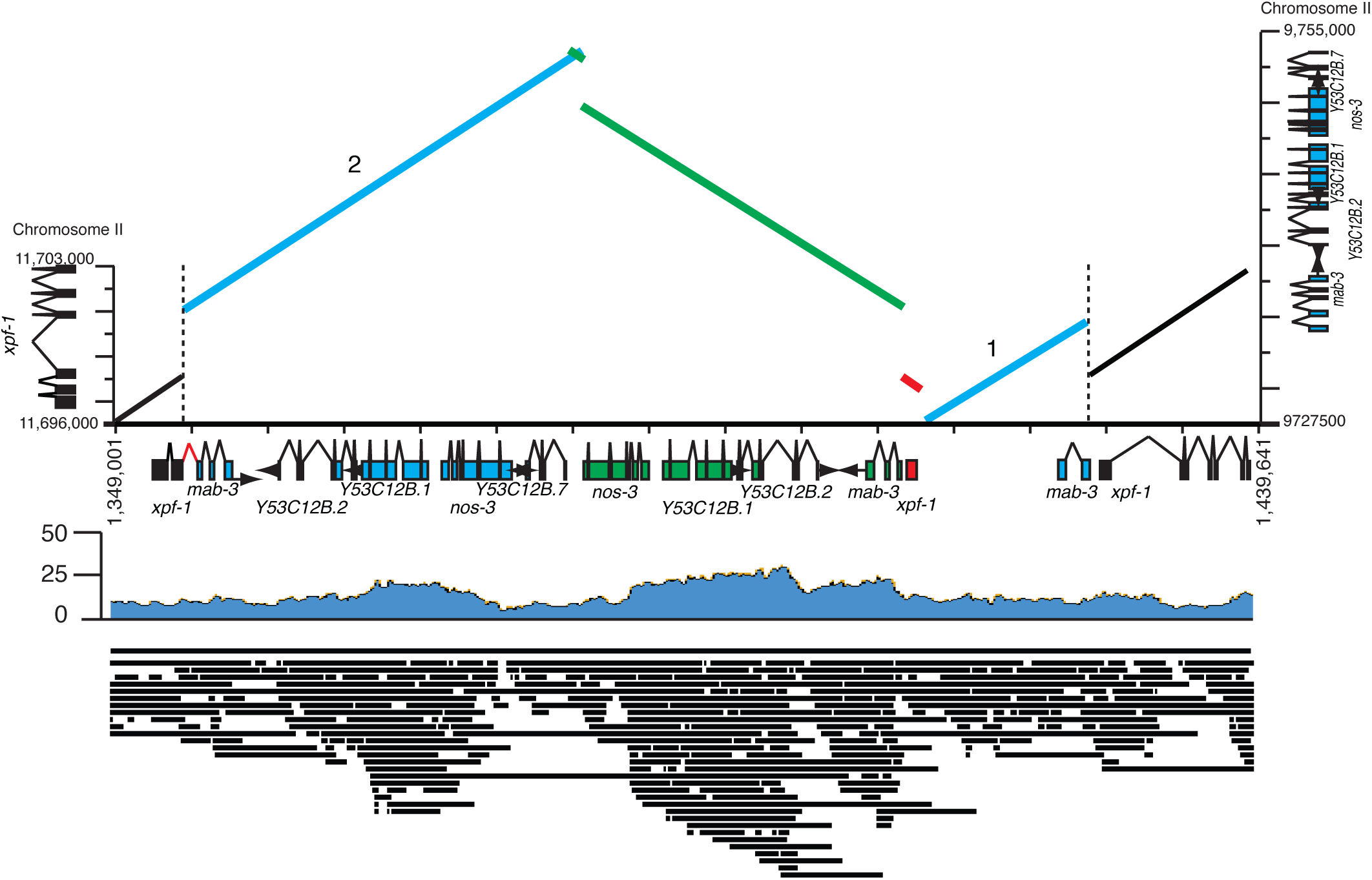
Schematic of the 40kb complex insertion of mab-3 region into the xpf-1 gene. The insertion is in contig 1766. The insertion is an inverted repeat.Shown below are selected MinION reads mapping to the region spanning the various breakpoints. Elucidating the structure of the complex *him-9(e1487)* acetaldehyde-induced mutation. (A) Schematic of the 40 kb insertion of the *mab-3* region into the *xpf-1* gene based on contig 1701. The insertion structure suggests a mutation event in which the insertion was duplicated to generate an inverted repeat of the *mab-3* region and the insertion point with *xpf-1* (in red). (B) The *mab-3* region with breakpoints denoted with arrows. Breakpoints detected by MinION are corroborated by Illumina read coverage and paired-end read alignment.

**Figure 4B.**
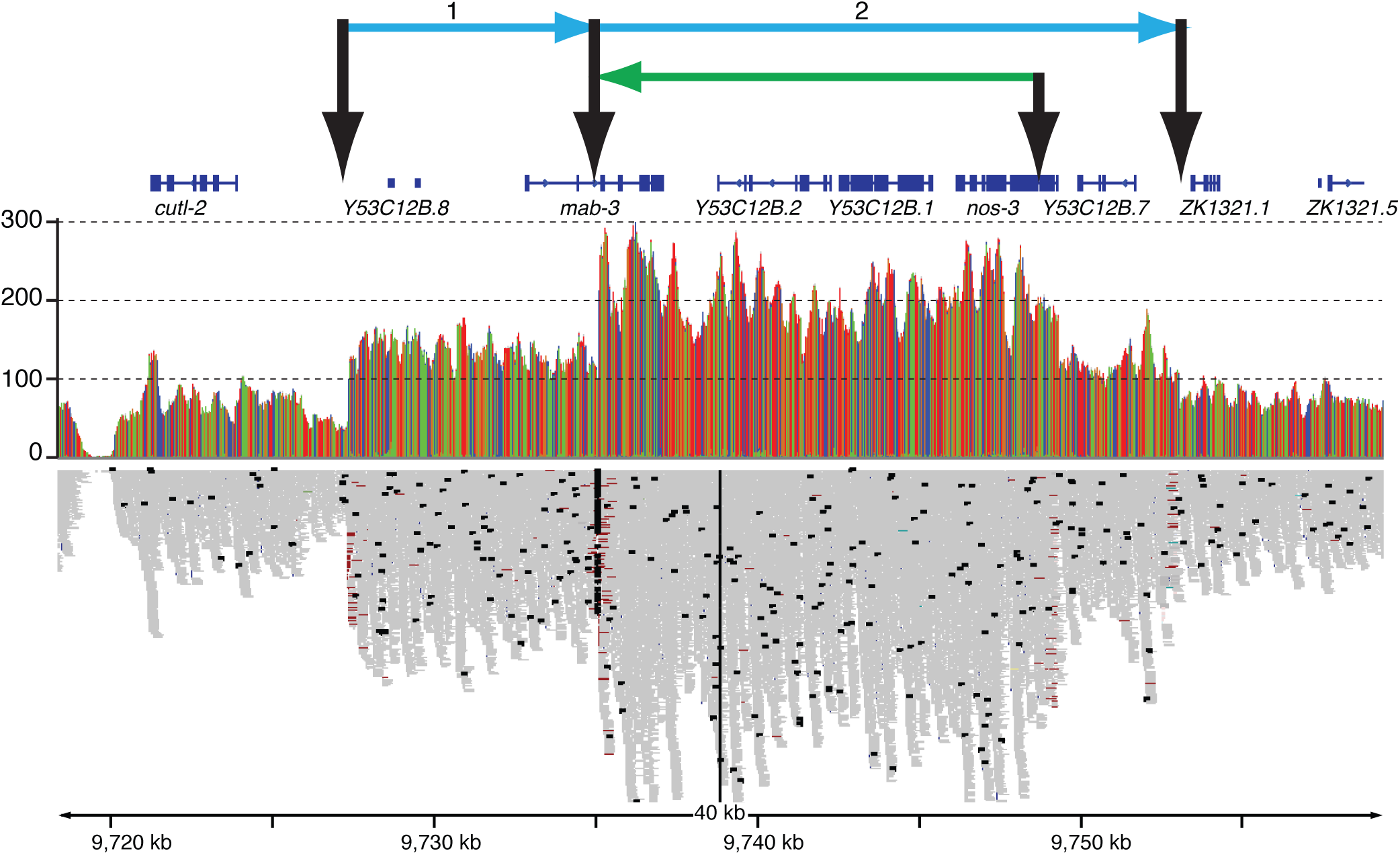
The mab-3 region showing breakpoints detecting in MinION sequencing data and Illumina read coverage.

The biolistic-mediated insertion was located on contig tig00000045, which aligned to the right arm of chromosome III consistent with the published location for *ruIs32* (Wormbase). From the MinION read assembly, it appears that the insertion contains three copies of the *Ppie-1::GFP::H2B::pie-1* transgene and two copies of the ampicillin gene from the plasmid and two partial copies of the *unc-119(+)* gene from the *unc-119* transgene. The structure of the insertion is complex, with the *unc-119(+)* sequence interspersed within the plasmid sequence suggesting a complex integration event (Figure 5A). Copy number changes and breakpoints in *pie-1* and *unc-119* were confirmed by the Illumina sequencing reads (Figure 5B). The integration event also appears to have generated a large duplication of approximately 2 Mb of DNA (chrIII:10,062,096-11,973,739) from the region near the insertion site (Figure 2). Given the position of the insertion, the wild type *unc-119* transgene should be genetically linked to the *unc-119(ed3)* mutation and should not be lost through outcrossing. Indeed, we were able to ascertain both the mutant and two wild type transgenic alleles of *unc-119* from the MinION contigs (Figure 5C).

**Figure 5.**
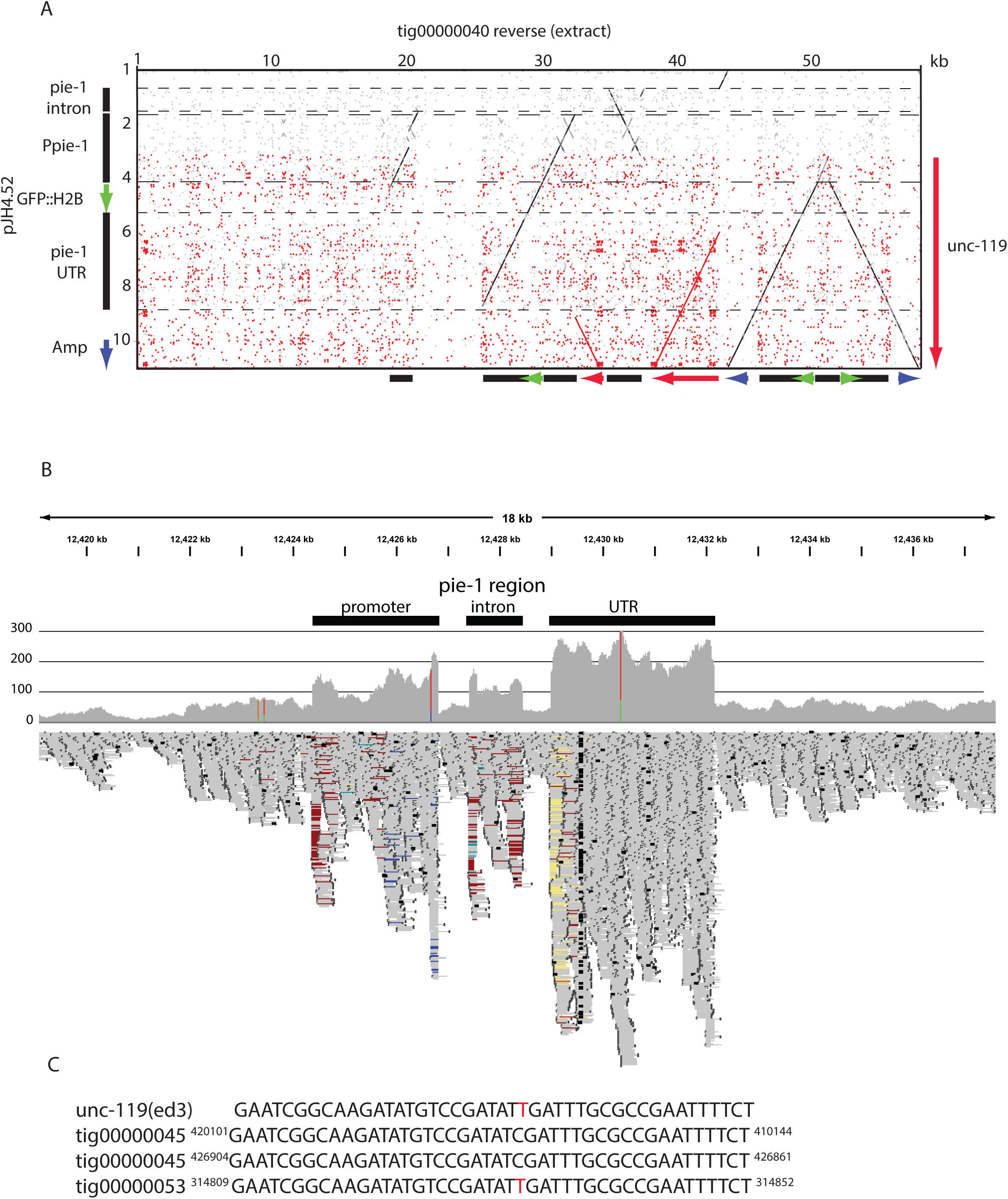
Elucidating the *ruIs32* insertion. (A) Dot plot of the contig 45 region that aligns with the pJH4.52 plasmid (pJH4.52 was used because pAZ132 sequence is not available. pAZ132 was derived from pJH4.52) and the *unc-119* gene. (B) Illumina read data illustrating the breakpoints and copy number changes identified in the MinION data. (C) The sequence of *unc-119* identified in the MinION data.

## Discussion

Advances in long read sequencing have opened new avenues for genome analysis. The relatively short sequencing reads produced by NGS allow for resequencing less complex, non-repetitive regions in well-characterized genomes. However, long sequence reads span repetitive regions and facilitate contiguous assemblies of large contigs. Similarly, long reads are essential for resolving complex chromosomal alterations such as those observed in tumour genomes. The major challenges previously limiting the use of long read TGS technology have been throughput, accuracy, and cost. The MinION sequencer has low equipment and consumable costs, and here we show that the MinION has achieved throughput and error rates that make it a viable option for the sequencing of novel genomes and genomes containing complex chromosomal rearrangements such as those observed in tumours. Continued advancements in both nanopore chemistry and basecalling will further improve throughput and accuracy. Additionally, with the advent of the high throughput ONT PromethION instrument genomes of larger sizes should be within easy reach.

We demonstrate here that the sequencing and *de novo* assembly of a large, low percent-GC (35.44), complex metazoan genome can be accomplished using MinION technology. We were able to assemble a near complete *C. elegans* genome with large contigs from less than 60-fold sequence coverage. MinION sequence read quality has increased to a level that 1D reads now achieve error rates <10% and result in more unique reads per flow cell.

Despite having higher error rates than NGS data, nanopore reads could be assembled into contigs with high concordance to the reference sequence. While consensus sequencing corrects for random sequencing errors, not all sequencing errors generated by MinION are random. Presently, MinION reads do not discern homopolymeric nucleotide tracts longer than 5 nucleotides. Consistent with this deficiency, the MinION assembled contigs did not contain any of the 396 >18 mer G-tracts known to be present in the *C. elegans* genome (Zhao et al. 2007). Pilon can improve the identity of homopolymeric tracts but not completely. Although homopolymeric runs were truncated in the MinION reads, the quality of sequence in their flanking regions was not affected. Illumina sequencing was more effective in sequencing homopolymeric tracts although we observed that Illumina reads were also affected by large G-tracts with lower read coverage in regions spanning G-tracts. Improvements in the ability to detect homopolymeric sequences from nanopore data are anticipated near-term.

For the purposes of this analysis, we assembled the genome using Canu alone and did not use manual finishing methods to bridge contigs. Assembled contigs were very large with an N50 contig size >1 Mb. A more contiguous genome assembly could be generated from the same data by using manual finishing approaches to bridge contigs. For example, it is clear from the LASTZ alignment that many of the contigs contain sequence overlaps between adjacent contigs. The genome assembly could be made more complete using long reads that extend from the ends of the contigs to find potential links between existing contigs resulting in a more contiguous genome assembly. The genome assembly could also be improved by changing the quality of the DNA sample before sequencing. Contig breakpoints were often in regions containing repetitive regions that were larger than the average read length. Increasing the number of reads longer than these repeat regions will facilitate the spanning of these regions. This could be accomplished by sequencing deeper or by preparing the genomic DNA to favour the formation of larger fragments.

Chromosome rearrangements are common in tumours and the identification of rearrangements is important for tumour characterization and treatment. Detecting chromosome rearrangements such as duplications, deletions and translocations pose challenges for sequencing-based approaches. NGS can detect copy number changes and identify potential breakpoints. However, complex rearrangements with multiple breakpoints and duplications can prove difficult to reassemble from NGS reads. We demonstrate that long MinION reads provide context for assembly of duplicated rearranged sequence and can delineate complex chromosome rearrangements. MinION long reads offer the opportunity to cheaply identify genome rearrangements in tumours including the highly complex chromothripsis events which result in thousands of clustered localized chromosomal rearrangements (Leibowitz et al. 2015).

Our sequencing and assembly of the *C. elegans* genome demonstrates the advancing capabilities of the MinION sequencer. Throughput and accuracy of the MinION platform continues to improve and is approaching 5-10 Gb of sequence per flowcell, which would generate sufficient sequence to assemble the 100 Mb *C. elegans* genome. The long read capabilities of MinION nanopore sequencing facilitate unambiguous assembly of chromosome structure, thereby eliminating the need for physical mapping. These properties allow for sequencing of new genomes or tumour genomes with structural chromosome changes. Combining MinION and Illumina sequencing currently can delineate the structure of novel genomes with higher base-level certainty. Alternative techniques, for example, using nanopore only event based correction methods (Loman et al. 2015), offer improved accuracies independent of hybrid NGS sequence correction. Combining this with future improvements in basecaller performance is anticipated to further remove the need for SBS in regard to high sequence accuracy.

## Acknowledgement

The authors would like to thank both Dr. Mark Akeson (MA) and Dr. Benedict Paten (BP) for supporting MJ and HEO, and for allowing use of the computer cluster to run assemblies and analyses. JRT and TPS thank the Canadian Institutes of Health Research (grant #10677) and Brain Canada Multi-Investigator Research Initiative Grant with matching support from Genome British Columbia, the Michael Smith Foundation for Health Research and the Koerner Foundation for funding. NJO and PH thank the Canadian Cancer Society (grant 702975). MJ and HEO thank the National Human Genome Research Institute of the US National Institutes of Health for funding their PI’s under award numbers HG006321 (MA), HG007827 (MA), and U54HG007990 (BP). We also thank Oxford Nanopore Technologies for access to hardware, software and sequencing chemistries.

